# Bispecific tau antibodies with additional binding to C1q or alpha-synuclein

**DOI:** 10.1101/2020.11.10.376301

**Authors:** Wim Hendricus Quint, Irena Matečko-Burmann, Irene Schilcher, Tina Löffler, Michael Schöll, Björn Marcus Burmann, Thomas Vogels

## Abstract

**Background:** Alzheimer’s disease (AD) and other tauopathies are neurodegenerative disorders characterized by cellular accumulation of aggregated tau protein. Tau pathology within these disorders is accompanied by chronic neuroinflammation, such as activation of the classical complement pathway by complement initiation factor C1q. Additionally, about half of the AD cases present with inclusions composed of aggregated alpha-synuclein called Lewy bodies. Lewy bodies in disorders such as Parkinson’s disease and Lewy body dementia also frequently occur together with tau pathology. Immunotherapy is currently the most promising treatment strategy for tauopathies. However, the presence of multiple pathological processes within tauopathies makes it desirable to simultaneously target more than one disease pathway.

**Methods:** Herein, we have developed three bispecific antibodies based on published antibody binding region sequences. One bispecific antibody binds to tau plus alpha-synuclein and two bispecific antibodies bind to tau plus C1q.

**Results:** The affinity of the bispecific antibodies to their targets compared to their monospecific counterparts ranged from nearly identical to one order of magnitude lower. All bispecific antibodies retained binding to aggregated protein in patient-derived brain sections. The bispecific antibodies also retained their ability to inhibit aggregation of recombinant tau, regardless of whether the tau binding sites were in IgG or scFv format. Mono- and bispecific antibodies inhibited cellular seeding induced by AD-derived pathological tau with similar efficacy. Finally, both Tau-C1q bispecific antibodies completely inhibited the classical complement pathway.

**Conclusion:** Bispecific antibodies that bind to multiple pathological targets may therefore present a promising approach to treat tauopathies and other neurodegenerative disorders.

## Introduction

Tauopathies are characterized by cellular accumulations of aggregated tau, such as the neurofibrillary tangles (NFTs) in Alzheimer’s disease (AD) [32, 77]. About 50% of the AD cases also contain α-synuclein (αSyn) inclusions called Lewy bodies [58, 67]. When present, Lewy body pathology contributes significantly to the clinical presentation of AD [23, 30, 40, 64, 70, 71, 76]. Likewise, tau pathology is often present in synucleinopathies such as Parkinson’s disease (PD) and dementia with Lewy bodies (DLB) [58, 67]. The presence of NFTs is not only strongly correlates with disease progression in tauopathies, but also contributes to neurodegeneration as well as clinical symptoms in different synucleinopathies [14, 15, 22, 36, 38, 85]. Furthermore, there is increasing evidence indicating a biological interaction between tau and αSyn pathology [5, 26, 29, 58, 68]. The last decade of research has also demonstrated many similarities in the pathological processes induced by tau and αSyn pathology: both proteins can progressively aggregate *via* templated misfolding (also known as ‘seeding’), accumulate into cells as fibrillar β-sheet rich aggregates, and can spread to other cells – thereby propagating their pathology [72, 77].

Tauopathy patients as well as transgenic animals with tau pathology also exhibit chronic neuroinflammation, which plays a key role in neurodegeneration [78]. One pro-inflammatory pathway that is robustly upregulated in response to tau pathology is the classical complement cascade [12, 19, 51, 82]. Since complement is also induced by AD-related Aβ plaque pathology [1, 55], it may be of particular importance to AD. The complement-initiating protein C1q is an extracellular protein that – in the context of neurodegeneration – is critical in synapse phagocytosis by microglia [19, 33], activation of neurotoxic A1 astrocytes [50], and neurodegeneration associated with activation of downstream complement components (e.g. C3, C5a) [13, 51, 82].

Immunotherapy is currently the most established approach for treating neurodegenerative disorders associated with protein aggregates (proteinopathies), such as tauopathies and synucleinopathies [73]. The dominating functional mechanism of these immunotherapies is antibody-mediated neutralization of misfolded and aggregated proteins in the extracellular space [84]. This process inhibits seeded aggregation in healthy cells, thereby potentially reducing the propagation of the disease progression [42]. Neutralizing antibodies against extracellular pro-inflammatory proteins have also been explored as a potential treatment strategy for proteinopathies [43, 66]. A neutralizing anti-C1q antibody inhibiting the classical complement pathway was shown to rescue both Aβ and tau pathology-induced synapse phagocytosis by microglia [19, 33].

There is growing consensus that the ultimately effective treatment for proteinopathies will consist of a combination treatment [65]. However, regulatory hurdles make it difficult to test combination treatments without showing efficacy of the individual drugs in a first stage. Furthermore, employing two drugs simultaneously complicates dose optimization in clinical trials. In the case of immunotherapy, combining two or more monoclonal antibodies may result in unsustainable treatment costs. Additionally, brain uptake of IgG is limited, and two monoclonal antibodies may compete for the same pathways, which may lead to lower uptake of both antibodies. This is particularly the case for the anticipated novel generation of antibodies, which bind to saturable proteins (e.g. transferrin) on the blood-brain barrier to promote their uptake in the brain parenchyma [35, 41].

The goal of the present study was to explore a novel approach: bispecific antibodies that simultaneously bind to two proteins involved in the pathological process of AD and other tauopathies. The use of bispecific antibodies that bind to two disease-related targets has increased in the past years in other fields (e.g. oncology), but – to the best of our knowledge – this has not yet been explored for the treatment of neurodegeneration [46]. In the present study we have developed bispecific antibodies based on published immunoglobulin G (IgG) complementarity-determining regions (CDRs) binding to tau plus αSyn as well as tau plus C1q. In addition, we show that the antibodies retained their target-binding and therapeutic effects in functional assays.

## Methods

### Human brain tissue

The brain samples were obtained from The Netherlands Brain Bank (NBB), Netherlands Institute for Neuroscience, Amsterdam (open access: www.brainbank.nl). All Material has been collected from donors for or from whom a written informed consent for a brain autopsy and the use of the material and clinical information for research purposes had been obtained by the NBB.

Paraffin-embedded middle frontal gyrus of a 68-year-old female patient with the Lewy body variant of AD (amyloid C, NFTs Braak VI, Lewy bodies Braak 6, APOE ε4/3) and the same region from a non-demented 69-year-old female (amyloid –, NFTs Braak I, Lewy bodies Braak 0, APOE ε3/3) were used for histological analysis. The middle frontal gyrus in this patient was previously characterized by the brain bank and shown to be heavily affected by Aβ plaques, neurofibrillary tangles, and Lewy body pathology. The *postmortem* delay was 3:30 hours for the patient tissue and 6:15 hours for the non-demented control tissue.

For the cellular seeding assay, a pool of frozen, non-fixed material was used. This consisted of angular gyrus of two male AD patients (ages 63 and 64 years old; 6; both APOE ε3/3) and superior parietal gyrus of four female AD patients (ages between 77 and 92 years old; APOE allele ε3/3, 3/4 and ε4/4). All samples were Braak stage VI for tau pathology and had a *post mortem* delay ranging from 3:00 to 4:45 hours.

### Production of bispecific antibodies

Production of bispecific antibodies was performed at Absolute Antibody (UK). The produced bispecific antibodies consist of a mouse IgG1 molecule with two scFv molecules fused at the carboxy-terminus of the heavy chain. Variable domains from publicly available DNA sequences were designed and optimized for expression in mammalian cells (HEK293) prior to being synthesized. The tau binding regions from mono- and bispecific variants of antibody A were derived from clone hu37D3-H9.v28 (US20190367592A1). The tau binding regions from mono- and bispecific variants of antibody B were derived from clone AB1 (WO2017005734A1) [16]. The αSyn binding regions were derived from clone M9E4 (US20200024336A1) [54]. The C1q binding regions were derived from clone M1 (US10590190B2) [33]. The sequences were subsequently cloned into Absolute Antibody cloning and expression vectors for mouse IgG1 and scFv. The C1q binding antibodies had an additional D265A mutation in the mouse IgG1 backbone to eliminate potential C1q binding in the Fc domain [6]. HEK293 cells were passaged to the optimum stage for transient transfection. Cells were transiently transfected with expression vectors and cultured for a further 6–14 days. An appropriate volume of cells was transfected with the aim to obtain 1–5 mg of purified antibody. Cultures were harvested and a one-step purification was performed by affinity chromatography. The antibodies were analyzed for purity by SDS-PAGE and the concentration was determined by UV spectroscopy. This format of the resulting bispecific antibodies is commonly referred to as IgG-scFv, IgG-scFv (HC) or BiS3.

### Enzyme-Linked Immuno Sorbent Assay (ELISA)

ELISA was performed at Absolute Antibody (UK). Maxisorb micro microplates were coated with 5μg/ml of antigen in PBS for 1 hour. Solutions were removed and plates were blocked overnight at 4°C in 1% casein solution. Solutions were removed and plates were washed once with PBS with 0.02% Tween-20. Antibody samples were added in duplicates and incubated for 1 hour with shaking at room temperature. Plates were washed 4 times with PBS supplemented with 0.02% Tween-20. Goat anti-mouse HRP conjugated secondary antibody (1:4000 dilution) was added and incubated for 1 hour with shaking at room temperature. Plates were washed 4 times with PBS with Tween-20 followed by washing twice with water. Detection was performed by incubation with the TMB substrate for 10 minutes, followed by 0.1M HCl. Absorbance was read out at 450nm.

### Purification of recombinant proteins

For the expression and purification of recombinant Tau40 (2N4R), *E. coli* BL21(λDE3) Star™ (Novagen) cells were transfected with a modified pET28b plasmid harboring full length Tau40 protein with an amino-terminal His-SUMO Tag (purchased *E. coli* codon optimized from GeneScript). Transformed cells were spread on LB-Agar plates containing Kanamycin (40μg/ml) (Sigma Aldrich) and grown overnight at 37°C. For protein expression and purification, the cells were pre-cultured overnight in 2xM9 medium [4] (supplemented with Kanamycin (40μg/ml) at 30°C. The main culture was inoculated with the precultured medium to an OD_600_ ≈ 0.1 and grown at 37°C until a cell density of OD_600_ ≈ 0.8 was reached at 37°C. Protein expression was induced by the addition of 1 mM isopropyl β-D-thiogalactoside (IPTG) (Thermo Scientific) for 16 h at 22°C. Cells were harvested by centrifugation at 5.000xg at 4°C for 20 min. The resulting cell pellet was washed once with ice cold PBS/EDTA buffer (0.137 M NaCl, 0.0027 M, 0.01 M Na_2_HPO_4_, 0.0018 M KH_2_PO_4_, 2mM EDTA), subsequently resuspended in 80 ml ice cold lysis/binding (0.025 M NaPi pH7.8, 0.5 M NaCl) buffer and lysed by four passes through an Emulsiflex (Avestin).

The cleared cell lysate was centrifuged at 22000xg for 1 hour at 4°C. The supernatant was directly applied to a HisTrap HP column (GE Healthcare), equilibrated with binding buffer, washed with 5 column volumes of the same buffer supplemented with 25 mM imidazole. Tau protein was eluted with a 150 mM imidazole step. Fractions containing Tau40 were pooled and dialyzed overnight, against 5L human SenP1 cleavage buffer (25mM TrisHCl, 150 mM NaCl, 1mM DTT, pH7.4). After dialysis SenP1 protease (Addgene plasmid #16356) [57] was added and the enzymatic cleavage was performed for 4 hours at room temperature. Separation of the His-SUMO-Tag and Tau40 was done by a second HisTrap HP column step and fractions containing cleaved Tau40 in the flow-through were collected, concentrated, and subsequently purified by gel filtration using a HiLoad 10/60 200pg column (GE Healthcare) pre-equilibrated with PBS buffer supplemented with 2mM EDTA. Pure tau fractions were concentrated to about 500 μM, flash frozen in liquid nitrogen, and stored at −80°C till usage.

Human αSyn was expressed from plasmid pRK172 (a kind gift of M. Goedert) [37] in *E. coli* BL21(λDE3) Star™ (Novagen) cells as described before [10, 34, 39]. Briefly, α-Syn was purified by a non-denaturing protocol by anion-exchange chromatography followed by a size-exclusion chromatography step using a Superdex75 Increase column (GE Healthcare). The α-Syn containing fractions were concentrated to about 500 μM and stored at −80°C.

### Bio-Layer Interferometry (BLI)

BLI experiments were performed on an OctetRED96 system (Fortébio) at 30°C. Recombinant Tau40, αSyn, and C1q (Abcam; ab96363) were biotinylated using the EZ-Link NHS-PEG4 Biotinylation Kit (Thermo Fisher Scientific) according to the manufacturer3’s instructions. Briefly, a biotin aliquot was freshly resolved in H_2_O, directly added to the protein solution to a final molar ratio of 1:1 in PBS buffer and the solution was gently mixed for 30 min at room temperature. Unreacted biotin was removed with Zeba Spin Desalting Columns (7 MWCO, Thermo Fisher Scientific). Biotin-labelled proteins were immobilized on the streptavidin (SA) biosensors (Fortébio) and the biosensors were subsequently blocked with EZ-Link Biocytin (Thermo Fisher Scientific). The different antibodies used were diluted and applied in a dose-dependent manner to the biosensors immobilized with the respective proteins. Experiments were performed in PBS buffer pH7.4 supplemented with 1% Bovine serum albumin (BSA) (Sigma-Aldrich) and 0.02% Tween (Fluka) to avoid non-specific interactions. Parallel experiments were performed for reference sensors with no antibodies bound and the signals were used for baseline subtraction during the subsequent data analysis. The association and dissociation periods were set to 300s and 500s, respectively. Data measurements and analysis were performed by using the Data acquisition 10.0 and the Data analysis HT 10.0 (Fortébio) software, respectively.

### Immunofluorescence

Immunofluorescent histological staining was based on a modified version of a previously published protocol [79]. All steps were performed at room temperature unless mentioned otherwise. Paraffin-embedded sections (8 μm) were deparaffinized and washed in PBS. Heat antigen retrieval was performed by immersing the sections in sodium citrate buffer (10 mM Sodium citrate, 0.05% Tween 20, pH 6.0) and boiling in a microwave for 10 minutes. The sections were then left to cool down at room temperature and subsequently incubated for 1 minute in TrueBlack lipofuscin autofluorescence quencher (Biotium, USA) diluted 1:20 in 70% ethanol. After washing, sections were blocked for 1 hour in horse serum and incubated in mono- and bispecific antibodies or commercial primary antibodies diluted in PBS overnight at 4°C. The following dilutions were used: experimental antibodies 1:2000 (from 1mg/ml stock), rabbit anti-Tau pS214 1:500 (ab170892, Abcam, UK), rabbit anti-αSyn 1:1000 (ab51253, Abcam, UK). The second day, sections were thoroughly washed and incubated for 1 hour in secondary antibodies anti-mouse IgG-Cy2, and anti-rabbit IgG-Cy3 (Thermo Fisher Scientific), both dissolved 1:1000 in PBS containing 10ug/ml Methoxy-X04 (Tocris Bioscience). After thoroughly washing, slides were coverslipped using anti-fade ProLong™ Gold Antifade Mountant (Thermo Fisher). Slides were imaged on a Zeiss LSM 700 confocal microscope.

### CH50-assay to measure classical complement

Human serum (Haemoscan, The Netherlands) was diluted 1/28 in dilution buffer and mixed with a previously reported complement-neutralizing dose of the C1q antibody: 1μg per test for the monospecific antibodies and 1.25μg for bispecific antibodies to have equal numbers of antibody molecules under all conditions [33]. The antibody:serum mixture was pre-incubated for 1 hour at 4°C, then mixed with bovine erythrocytes at 37°C according to the manufacturer’s instructions (Haemoscan, The Netherlands). Stop solution was applied after 30 minutes, samples were centrifuged for 10 minutes at 400xg and subsequently measured at OD_415_ to determine the amount of cell lysis – according to the manufacturer’s instructions (Haemoscan, The Netherlands). IgG containing samples were measured in duplicates and data was pooled from 5 independently prepared experiments. Dilution buffer without serum was used as a negative control and serum without experimental antibodies was used as a positive control and the resulting OD_415_ values were averaged to determine 100% hemolysis.

### Cell-free tau aggregation assay

Recombinant Tau441 (2N4R) P301L (Analytik Jena, T-1014-1) at 1 μM final concentration, was incubated with 30 μM sodium octadecylsulfate (ODS) and 1 μM heparin in reagent buffer (20 μM Thioflavin T, 5 mM 1,4-dithioerythreitol, 100 mM NaCl, 10 mM HEPES pH 7.4) for 15 hours at 37°C in black no-binding 96 well plates. IgG and IgG-scFv were used in the same concentration as recombinant tau (1 μM). Compounds were incubated with the before mentioned tau-Heparin-ODS-Buffer solution. Six technical replicates were performed. Immediately after preparation a baseline measurement was carried out and following 4 and 15 hours of incubation at 37°C, fluorescence was again detected by using 450 nm excitation and 485 nm emission.

### Extraction of sarkosyl insoluble tau

The preparation of sarkosyl insoluble brain fraction was performed as described previously [21]. AD brain tissue was homogenized in 3 volumes (v/w) of cold H buffer (10 mM Tris, 1 mM EGTA, 0.8 M NaCl, 10% sucrose, pH 7.4, containing 1 mM PMSF) with protease inhibitor (EMD Millipore, 539131). After 20 minutes incubation on ice the homogenate was spun at 27.200×g for 20 minutes at 4°C. Supernatants were supplemented with 1% sarkosyl and 1% 2-mercaptoethanol final concentration and incubated for 1 hour at 37°C on orbital shaker. The samples were then centrifuged at 150.000×g for 35 minutes at room temperature. The pellet was resuspended in TBS (10 mM Tris, 154 mM NaCl) and the insoluble fraction was used as described below.

### Capillary electrophoresis-based immunoassay

Automated separation and immunostaining of tau was carried out using a capillary-based immunoassay, WES™ (proteinsimple^®^). Insoluble sarkosyl extraction samples (0.2 mg/mL) before and after sonication for 2 minutes were applied to a 25-capillary cartridge with a 2 to 440 kDa matrix, according to the manufacturer’s protocol. After samples and antibody (Tau-13, BioLegend Inc.) have been pipetted into the pre-filled assay plate purchased from the manufacturer, sample loading, separation, immunoprobing, washing and detection were performed automatically by WES™ Western system. Quantitative data analysis was performed with Compass for SW software (Bio-Techne). The areas under the curve were determined for the subsequent analysis.

### Cellular seeding assays with AD-derived tau

SH-SY5Y-hTau441 P301L cells were kept in culture medium (DMEM medium, 10% FCS, 1% NEAA, 1% L-Glutamine, 100 μg/ml Gentamycin, 300 μg/ml Geneticin G-418) for ~2 days until 80–90% confluency was reached. Next, cells were differentiated in culture medium supplemented with 10 μM retinoic acid for 5 days changing medium every 2 to 3 days. Differentiated SH-SY5Y-hTau441 P301L cells were incubated with sarkosyl extracts from brain in combination with two different concentrations of the antibodies. Therefore, 2.5 μg total protein of brain extracts were mixed with 300 nM or 30 nM of the antibodies in Opti-MEM and incubated overnight at 4°C. On the same day, SH-SY5Y-hTau441 P301L cells were seeded in culture medium on 96-well plates at a cell density of 5 × 10^4^ cells/well. On the next day, the tau-antibody mixtures were incubated for 10 minutes with Lipofectamine 2000 (Invitrogen) in Opti-MEM, followed by adding these mixtures to the cells and incubation for 48 hours at 37°C. Two days after tau treatment, cells were harvested. To this end, cells were washed once with cold PBS and harvested in 50 μL FRET lysis buffer (Cisbio) per well and analyzed according to the manufacturer’s protocol.

Briefly, samples were diluted 1:2 in lysis buffer and the Anti-human TAU-d2 conjugate as well as the Anti-human Tau-Tb^3+^-Cryptate conjugate were diluted 1:50 in diluent solution and premixed. Thereafter, 16 μL of the lysates and 4 μL premixed conjugates were applied to a white 396 well plate and incubated approximately 20 hours at RT on a shaker. Fluorescence emission at two different wavelengths (665 nm and 620 nm) was performed on a multilabel plate counter (Victor 3V, PerkinElmer). The signal ratio was calculated using the following formula: (Signal 665 nm / Signal 620 nm) × 10^4^.

### Statistical analyses

Statistical analyses and visualization of results was performed in Prism 8 (Graphpad Software, San Diego, CA, USA). Experimental antibodies were compared to vehicle control and each other using a Welch test with correction for multiple comparisons using the Benjamini-Hochberg procedure to keep the false discovery rate below 0.05 (cell free tau aggregation assay; CH50 assay). This test assumes normally distributed data, but no equal variances between different conditions. For the cellular tau seeding assay, we used the non-parametric Kruskal-Wallis test with Dunn’s multiple comparisons test, which does not assume normally distributed data. The threshold of corrected P-values in our analysis is 0.05. Individual data are presented in the graphs along with means and 95% confidence intervals - unless stated otherwise in the figure legends. Comparisons between monospecific antibodies and their bispecific counterparts were indicated in the graphs with an asterisk if the corrected P-value was below the threshold and denoted as no difference (n.d.) when the corrected P-value did not cross the threshold.

## Results

### Bispecific tau antibodies with additional binding to αSyn and C1q

To validate a novel approach to immunotherapy for tauopathies, we have developed and characterized three bispecific antibodies (Table 1). Antibody A binds to tau plus αSyn, with two tau bindings sites attached as scFv to the anti-αSyn IgG (Figure 1A, Table 1). Antibody B-I is an anti-tau IgG fused to two C1q-binding scFvs (Figure 1B, Table 1). Antibody B-II has the same binding regions as antibody B-II, except that this antibody is an anti-C1q IgG fused to two tau-binding scFvs (Figure 1C, Table 1). Antibody A allowed us to study immunotherapy binding to two proteinopathy-related targets. Antibody B-I and B-II allowed us to study immunotherapy to tau an inflammation-related target. Furthermore, since antibodies B-I and B-II were identical except for their formats, this also allowed us to compare the influence of the bispecific antibody format on functionality. Antibodies were compared to their monospecific counterparts. Successful production of bispecific antibodies was confirmed by SDS-Page. All three blots looked similar with a single band at ~198kDa under non-denaturing conditions (Figure 1D, 1E, 1F). This is consistent with an IgG molecule having a molecular weight of ~150kDa plus two scFv molecules with a MW of ~25kDa each.

**Table 1.**
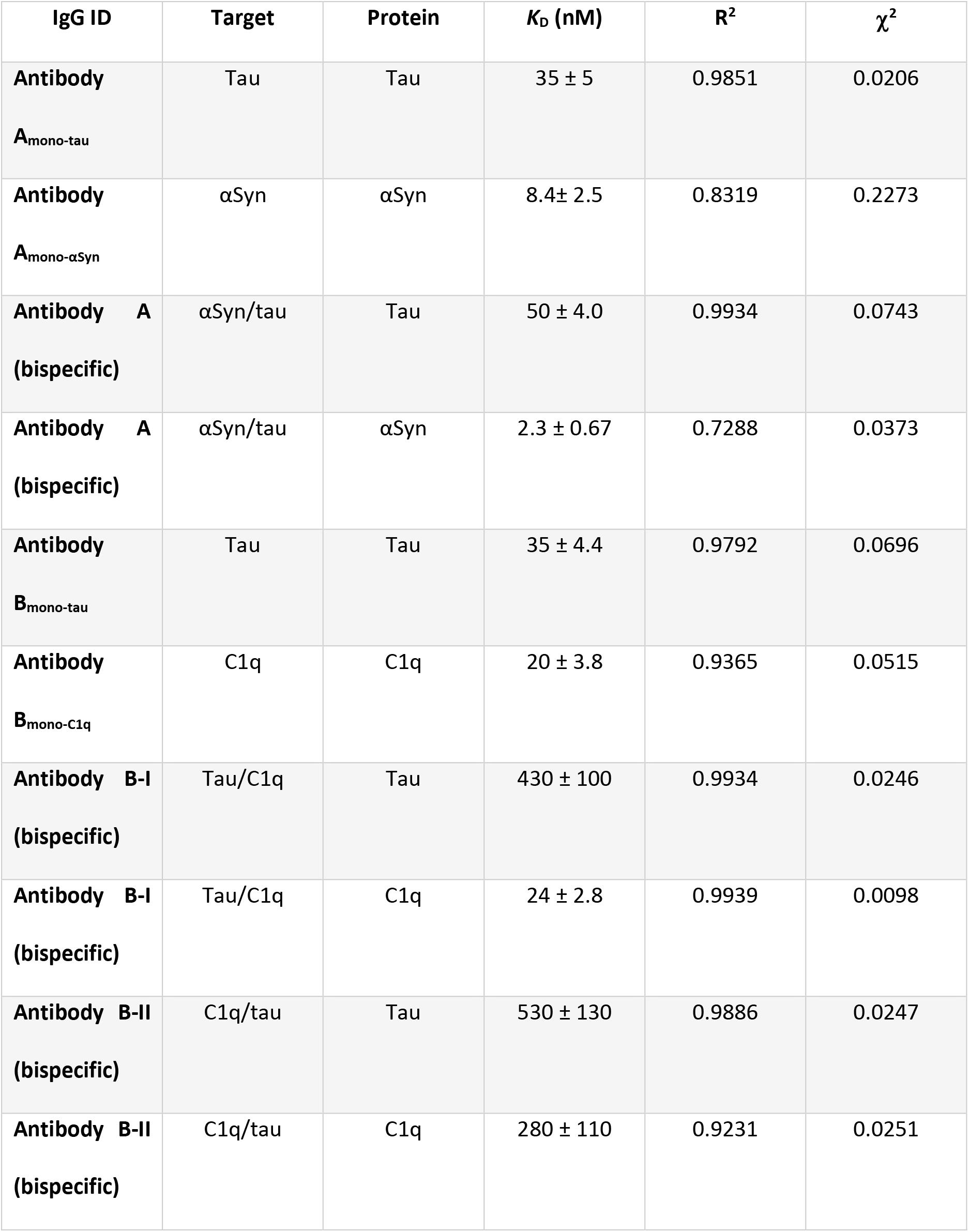
Antibody binding characteristics.

| **IgG ID** | **Target** | **Protein** | **$K_D$ (nM)** | **$R^2$** | **$\chi^2$** |
| --- | --- | --- | --- | --- | --- |
| **Antibody**<br>**A<sub>mono-tau</sub>** | Tau | Tau | $35 \pm 5$ | 0.9851 | 0.0206 |
| **Antibody**<br>**A<sub>mono-αSyn</sub>** | αSyn | αSyn | $8.4 \pm 2.5$ | 0.8319 | 0.2273 |
| **Antibody A**<br>**(bispecific)** | αSyn/tau | Tau | $50 \pm 4.0$ | 0.9934 | 0.0743 |
| **Antibody A**<br>**(bispecific)** | αSyn/tau | αSyn | $2.3 \pm 0.67$ | 0.7288 | 0.0373 |
| **Antibody**<br>**B<sub>mono-tau</sub>** | Tau | Tau | $35 \pm 4.4$ | 0.9792 | 0.0696 |
| **Antibody**<br>**B<sub>mono-C1q</sub>** | C1q | C1q | $20 \pm 3.8$ | 0.9365 | 0.0515 |
| **Antibody B-I**<br>**(bispecific)** | Tau/C1q | Tau | $430 \pm 100$ | 0.9934 | 0.0246 |
| **Antibody B-I**<br>**(bispecific)** | Tau/C1q | C1q | $24 \pm 2.8$ | 0.9939 | 0.0098 |
| **Antibody B-II**<br>**(bispecific)** | C1q/tau | Tau | $530 \pm 130$ | 0.9886 | 0.0247 |
| **Antibody B-II**<br>**(bispecific)** | C1q/tau | C1q | $280 \pm 110$ | 0.9231 | 0.0251 |

**Figure 1.**
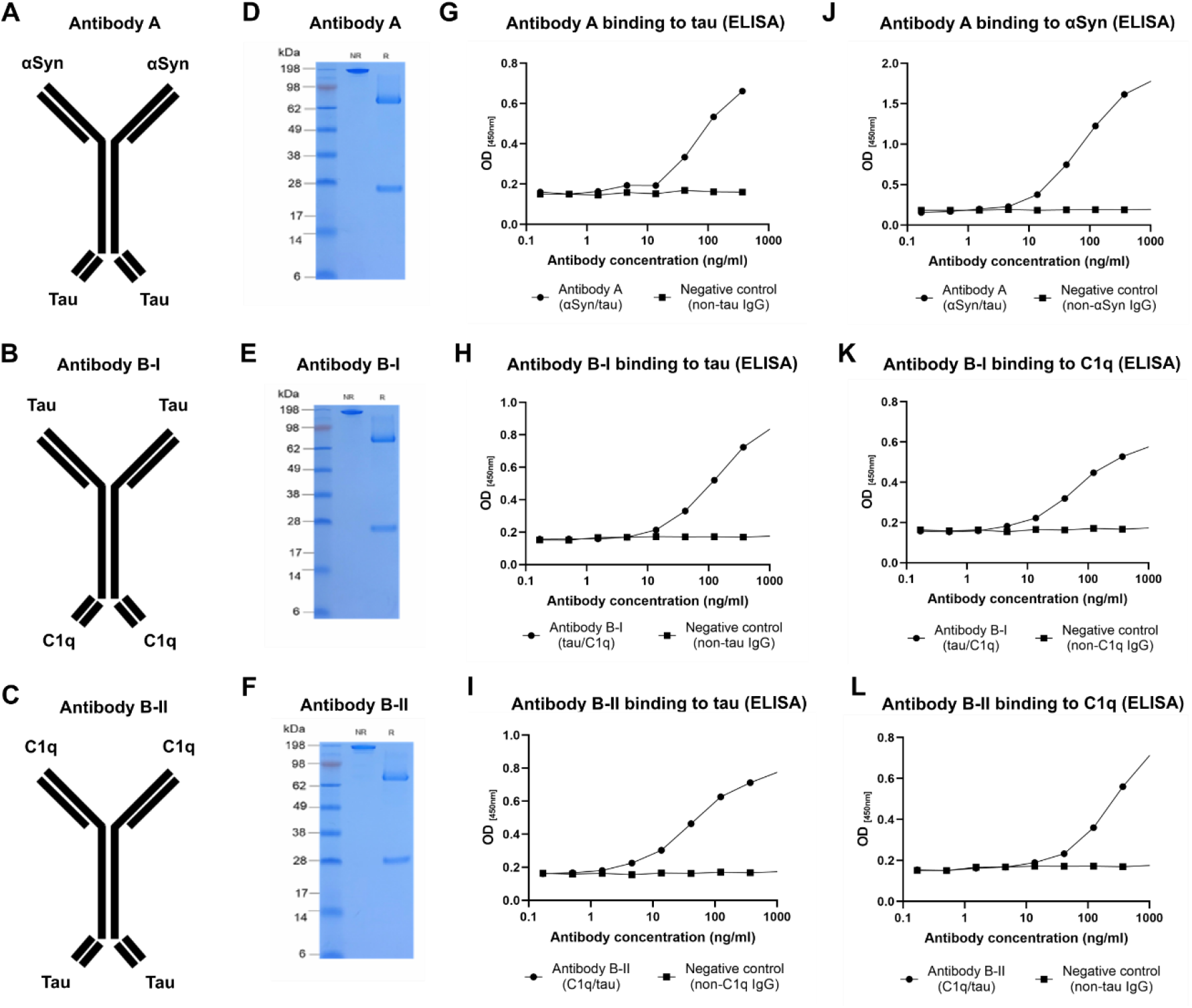
Antibody characteristics. **(A)** Antibody A is an anti-αSyn IgG with two additional tau-binding scFv domains fused to the Fc domain. **(B)** Antibody B-I is anti-tau IgG with two additional C1q-binding scFv domains fused to the Fc domain. **(C)** Antibody B-II is anti-C1q IgG with two additional tau-binding scFv domains fused to the Fc domain. **(D)** SDS-PAGE showing the molecular weight of Antibody A under non-reducing conditions in the middle lane and under reducing conditions in the right lane. **(E)** SDS-PAGE showing the molecular weight of Antibody B-I under non-reducing conditions in the middle lane and under reducing conditions in the right lane. **(F)** SDS-PAGE showing the molecular weight of Antibody B-II under non-reducing conditions in the middle lane and under reducing conditions in the right lane. **(G)** ELISA showing binding of Antibody A to recombinant tau. **(H)** ELISA showing binding of Antibody B-I to recombinant tau. **(I)** ELISA showing binding of Antibody B-II to recombinant tau. **(J)** ELISA showing binding of Antibody A to recombinant αSyn. **(K)** ELISA showing binding of Antibody B-I to recombinant C1q. **(L)** ELISA showing binding of Antibody B-II to recombinant C1q.

### Confirmation of target-binding and binding kinetics

To confirm that the bispecific antibodies retained binding to their targets, we tested them with ELISA. Antibody A efficiently recognized both tau and αSyn (Figure 1G, 1J). Both antibody B-I and antibody B-II recognized tau and C1q (Figure 1H, 1I, 1K, 1L). We then used Octet BLI to determine the binding affinities of the bispecific antibodies, which are summarized in Table 1. Monospecific antibody A_mono-tau_ and bispecific antibody A bound with similar affinity to tau (*K*_D_ values of 35nM and 50nM, respectively) (Table 1, Figure 2A). Similarly, monospecific antibody A_mono-αSyn_ and bispecific antibody A bound with similar affinity to αSyn (*K*_D_ values of 8.4nM and 2.3nM, respectively) (Table 1, Figure 2B). However, monospecific antibody B_mono-tau_ had an approximately 10-fold higher affinity compared to both bispecific antibodies B-I and B-II (*K*_D_ value of 35nM compared to 430nM and 530nM, respectively) (Table 1, Figure 2C, 2E). Antibody B_mono-C1q_ also had an ~10-fold higher affinity to C1q compared to the bispecific antibody B-I (*K*_D_ value of 20nM compared to 240nM) (Table 1, Figure 2D).

**Figure 2.**
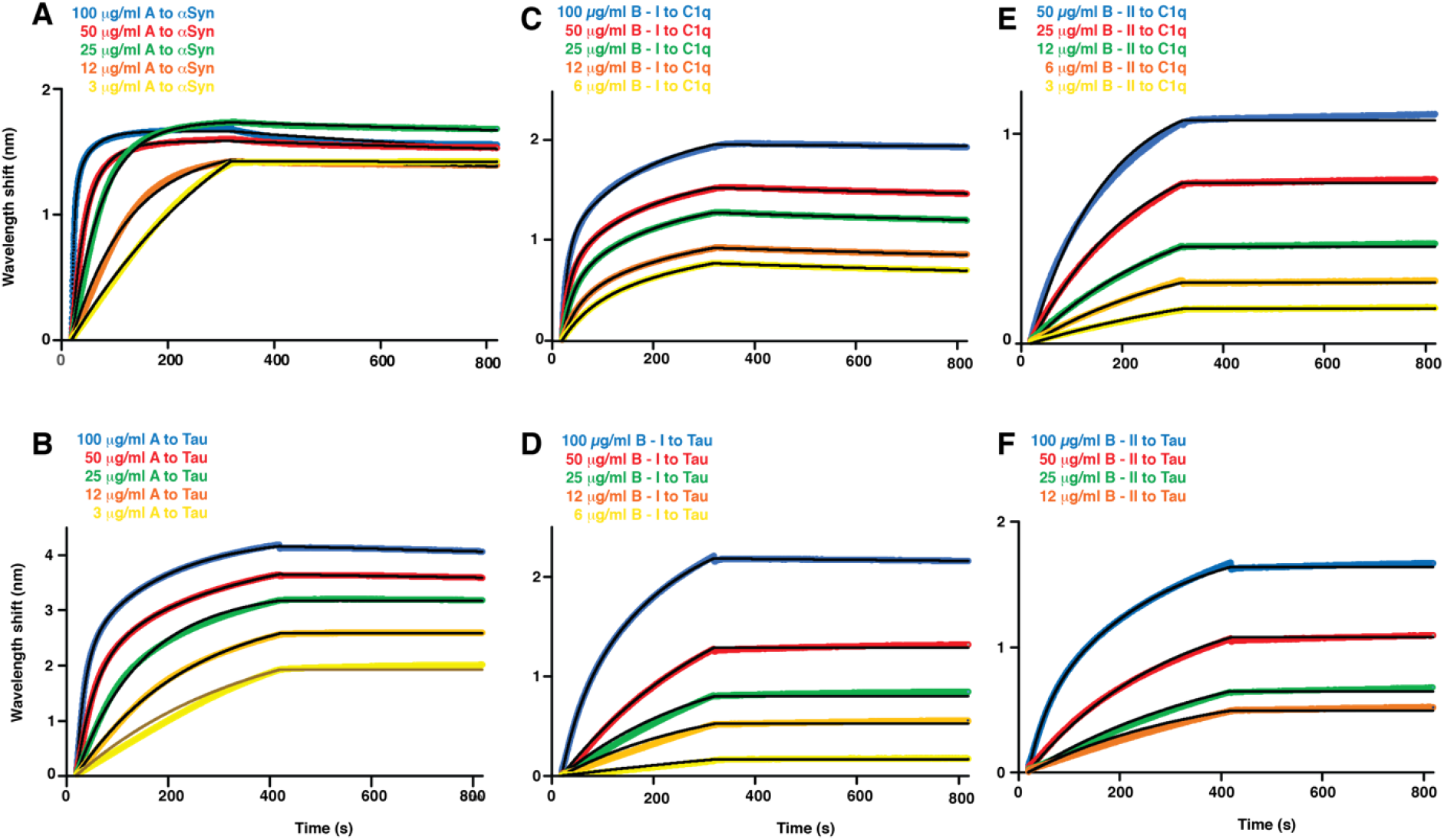
Sensograms of different bispecific antibodies binding to respective antigens using streptavidin (SA) sensors on an Octet Red96. Tau40, C1q, and αSyn were biotinylated and subsequently immobilized to the sensor. Antibodies were applied in a dose-dependent manner as indicated. (**A**) Antibody A to αSyn. (**B**) Antibody A to Tau40. (**C**) Antibody B-I to C1q. (**D**) Antibody B-I to Tau40. (**E**) Antibody B-II to C1q. (**F**) Antibody B-II to Tau40.

### Binding to neuropathology in patients

To determine if the bispecific antibodies were able to recognize NFTs and Lewy bodies, we used them as primary antibodies to stain human brain sections of an AD patient with Lewy bodies. Sections were co-labelled with Methoxy-X04, a Congo Red-derivative recognizing β-sheet–rich structures like amyloid plaques, NFTs, and Lewy bodies [44]. When tau binding sites were examined, sections were additionally labelled with an antibody that recognizes tau phosphorylated at the AD-related site serine 214 (pS214). When αSyn binding sites were examined, sections were additionally labelled with an antibody which recognizes αSyn phosphorylated at the synucleinopathy-related site serine 129 (pS129). Triple-positive structures with typical NFT or Lewy body morphology were therefore interpreted as specific binding. We examined paraffin embedded middle frontal gyrus sections of an AD patient with Lewy bodies. The slices contained abundant amyloid plaque, tau, and αSyn pathology. Antibody A recognized NFTs in the patient brain, but not in the control brain (Figure 3A). The same antibody also bound Lewy bodies in patient brain, but not in control brain (Figure 3B). NFTs were also detected with Antibody B-I (Figure 3C) and antibody B-II (Figure 3D). Although C1q was reported to be detectable around Thioflavin-positive plaques [1], we could not obtain a specific signal using this assay that could be reliably interpreted (results not shown).

**Figure 3.**
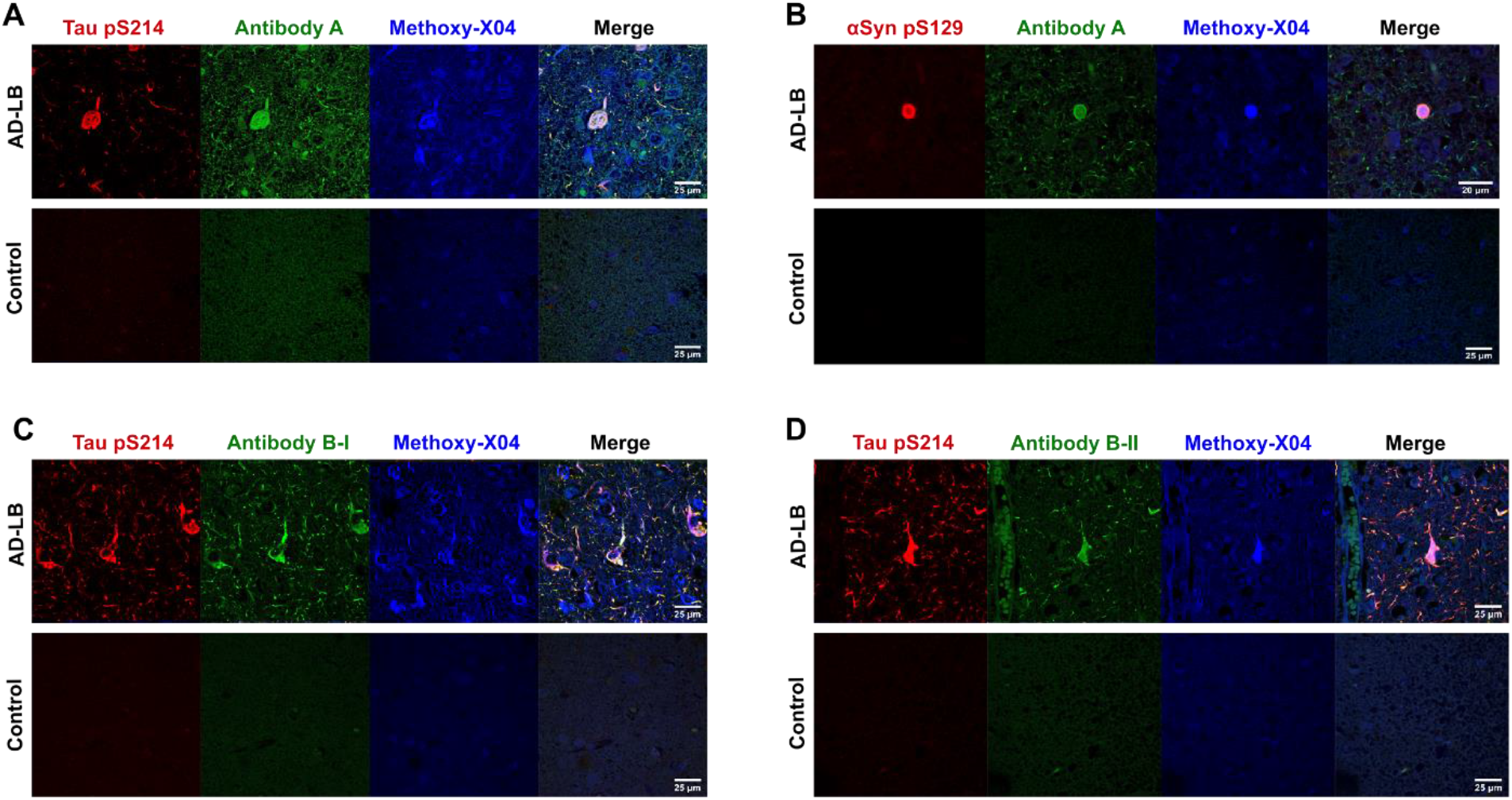
Histological detection of neurofibrillary tangles and Lewy bodies in patient brain sections. Bispecific antibodies were tested for their ability to recognize proteinopathy in an AD patient with Lewy bodies. Slides were co-labelled with β-sheet–specific dye Methoxy-04 and tau pS214 or αSyn pS129. **(A)** Antibody A recognized pS214 and Methoxy-X04 positive neurofibrillary tangles in patient tissue. Signal was absent in control. **(B)** Antibody A recognized pS129 and Methoxy-X04 positive Lewy bodies in patient tissue. Signal was absent in control. **(C)** Antibody B-I recognized pS214 and Methoxy-X04 positive neurofibrillary tangles in patient tissue. Signal was absent in control. **(D)** Antibody B-II recognized pS214 and Methoxy-X04 positive neurofibrillary tangles in patient tissue. Signal was absent in control.

### Inhibition of de novo aggregation of tau

Since tau is the common target of all three bispecific antibodies, we focused most of our analysis on functional assays related to tau pathology. We used a modified version of a previously established cell-free aggregation assay (Figure 4A). Similar assays were previously used to test the ability of tau antibodies to inhibit aggregation [2, 3, 17, 45]. Antibodies were co-incubated with tau protein (2N4R) and heparin was subsequently added to start the aggregation process. β-sheet binding dye thioflavin-T was used to estimate the presence of aggregates. Indeed, we observed robust aggregation without anti-tau antibodies, which was absent when tau was omitted (Figure 4B). The mean values of these conditions were used to determine 0% and 100% aggregation for normalization.

**Figure 4.**
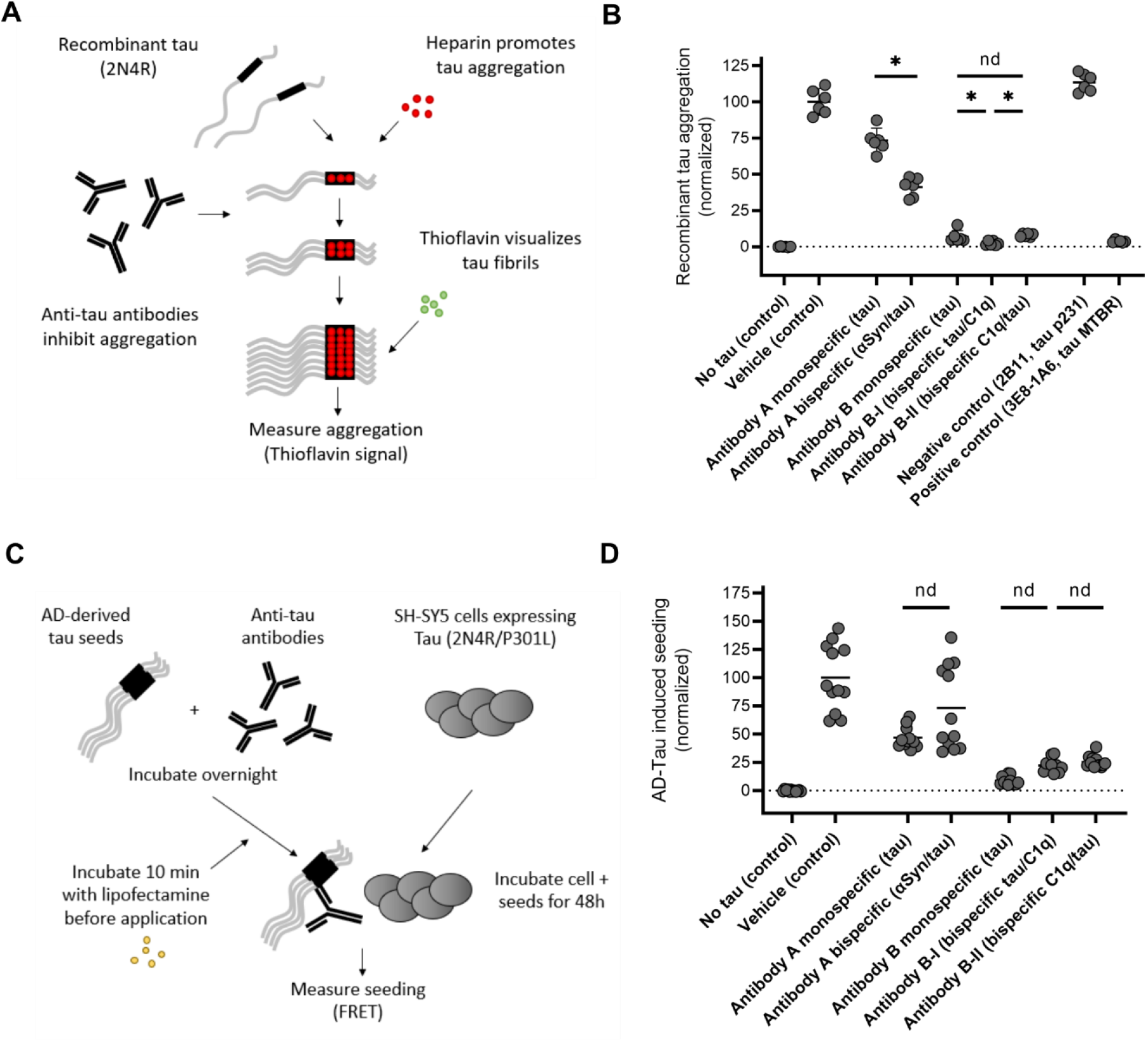
Comparison of mono-and bispecific antibodies in their ability to inhibit tau aggregation and tau seeding. (**A**) Schematic representation of the recombinant tau-based aggregation assay. (**B**) Comparison of antibodies in recombinant tau-based aggregation assay. Horizontal lines represent the mean and error bars represent the 95% confidence intervals. Only comparisons between monospecific antibodies and their bispecific counterparts are highlighted here. Detailed information is described in the corresponding results section. (**C**) Schematic representation of the AD tau-based cellular seeding assay. (**D**) Comparison of antibodies in recombinant tau-based cellular seeding assay. Horizontal lines represent the mean. No confidence intervals are shown because the data did not follow a Gaussian distribution. Only comparisons between monospecific antibodies and their bispecific counterparts are highlighted here. Detailed information is described in the corresponding results section.

Antibody A_mono-tau_, which binds to the amino-terminal of tau, reduced aggregation of recombinant tau to a mean value of 73.3% (corrected p= <0.001, 95% confidence intervals 64.7–81.8). Its bispecific counterpart, Antibody A, reduced aggregation to 41.2% (corrected p= <0.001, CI 34.2–48.1). Surprisingly, bispecific Antibody A was more effective at reducing aggregation compared to its monospecific counterpart (corrected p= <0.001) (Figure 4B). These results indicate that for bispecific antibodies in this format targeting the amino-terminus, it may be more efficacious to have anti-tau scFvs rather than anti-tau IgG in this assay.

To test the influence of the bispecific antibody format further, we tested both antibody B-I and B-II and their monospecific counterpart in the same assay (Figure 4B). The bispecific antibodies behave similar to one another, except that their IgG and scFv binding sites are in opposite configuration. The tau binding regions of the antibodies recognize the mid-domain of tau, partly overlapping with the first repeat domain [16]. Antibody B_mono-tau_ inhibited tau aggregation to only 6.9% (corrected p= <0.001, CI 2.6–11.2). Bispecific antibody B-I inhibited tau aggregation to 2.4% (corrected p= <0.001, CI 0.9–4.0). Antibody B_mono-tau_ inhibited tau aggregation to 8.0% (corrected p= <0.001, CI 6.8–9.2). Antibody B-I was slightly more effective at inhibiting tau aggregation compared to Antibody B_mono-tau_ and antibody B-II (both corrected p-values below 0.001).

Antibody B_mono-tau_ was more effective than Antibody A_mono-tau_ despite similar affinity to tau (corrected p= <0.001) (Figure 2B, Table 1), indicating that mid-domain antibodies are more effective at inhibiting recombinant tau aggregation than amino-terminal antibodies. Surprisingly, Antibody B_mono-tau_ was as effective at inhibiting tau aggregation as the positive control antibody 3E8-1A6 (corrected p= 0.116), which inhibited tau aggregation to only 3.7% compared to vehicle control (corrected p= <0.001, CI 2.8– 4.6). Antibody 3E8-1A6 binds to one of the two hexapeptides in the repeat domain of tau, which are supposed to be responsible for aggregation [7, 8]. This antibody was selected because antibodies targeting this domain were previously shown to potently inhibit recombinant tau aggregation in similar assays [45, 63]. Negative control antibody 2B11, which only recognizes tau phosphorylated at threonine 231 (absent on recombinant tau), did not reduce tau aggregation (Figure 4B)

### Inhibition of cellular seeding induced by pathological tau from AD brain

Cellular uptake of pathological tau from the extracellular space can lead to seeded aggregation of physiological tau *in vitro*. This process can be blocked by pre-incubation of pathological tau with antibodies and similar assays have been used previously to identify efficacious tau antibodies [16, 60, 61, 74, 84]. We developed a similar assay to test if the bispecific antibodies retained their ability to inhibit seeded aggregation induced by AD-derived sarkosyl-insoluble tau (Figure 4C). Seeding was detected using a sensitive FRET assay as previously described [16]. After estimating the tau concentration of the insoluble fraction of our AD brain extract, we tested a range of doses of both AD-Tau and lipofectamine. We determined that 2.5 μg AD-Tau with 1μL lipofectamine led to robust seeding without negative effects on cell-viability. In a next step, we tried a range of doses of commercial anti-human tau antibody HT7, which was previously used as a positive control in similar assays [59, 60, 69, 74] (not shown). Based on these results and the dosage used in previous publications [16, 74], we determined 300nM to be a suitable neutralizing concentration and this dose was used for further experiments.

We next tested the bispecific antibodies and their monospecific counterparts in this assay. High variability and lack of Gaussian distribution were observed in the vehicle control condition but not in most antibody conditions, leading us to use a relatively strict non-parametric statistical test (see Methods section). Antibody A_mono-tau_ and Antibody A both bind to the amino-terminal of tau. Antibodies against this domain were previously shown to be not very effective at reducing seeding with of sarkosyl insoluble AD brain extract [16, 74]. Indeed, antibody A_mono-tau_ did not reduce cellular seeding (correct p= 0.901, mean value of 46.9%, CI 40.6–53.2). Likewise, Antibody A also did not reduce cellular seeding (corrected p= >0.999, mean value of 73.2%, CI 49.4–97.0). No difference was observed between Antibody A_mono-tau_ and Antibody A in this assay (corrected p= > 0.999). In contrast, pre-incubation with Antibody B_mono-tau_ neutralized AD-Tau-induced seeding to 8.9% (corrected p= <0.001, CI 6.5–11.3) Antibody B-I also inhibited seeding (corrected p= <0.001, mean value of 22.3%, CI 17.9–26.7), as did Antibody B-II (corrected p= <0.001, 25.4%, CI 22.2-28.7) (Figure 5B). No differences were observed between Antibody B_mono-tau_ and Antibody B-I (corrected p= >0.999) or antibody B-II (corrected p= 0.318). Likewise, no differences were observed between bispecific antibodies B-I and B-II (corrected p= >0.999).

**Figure 5.**
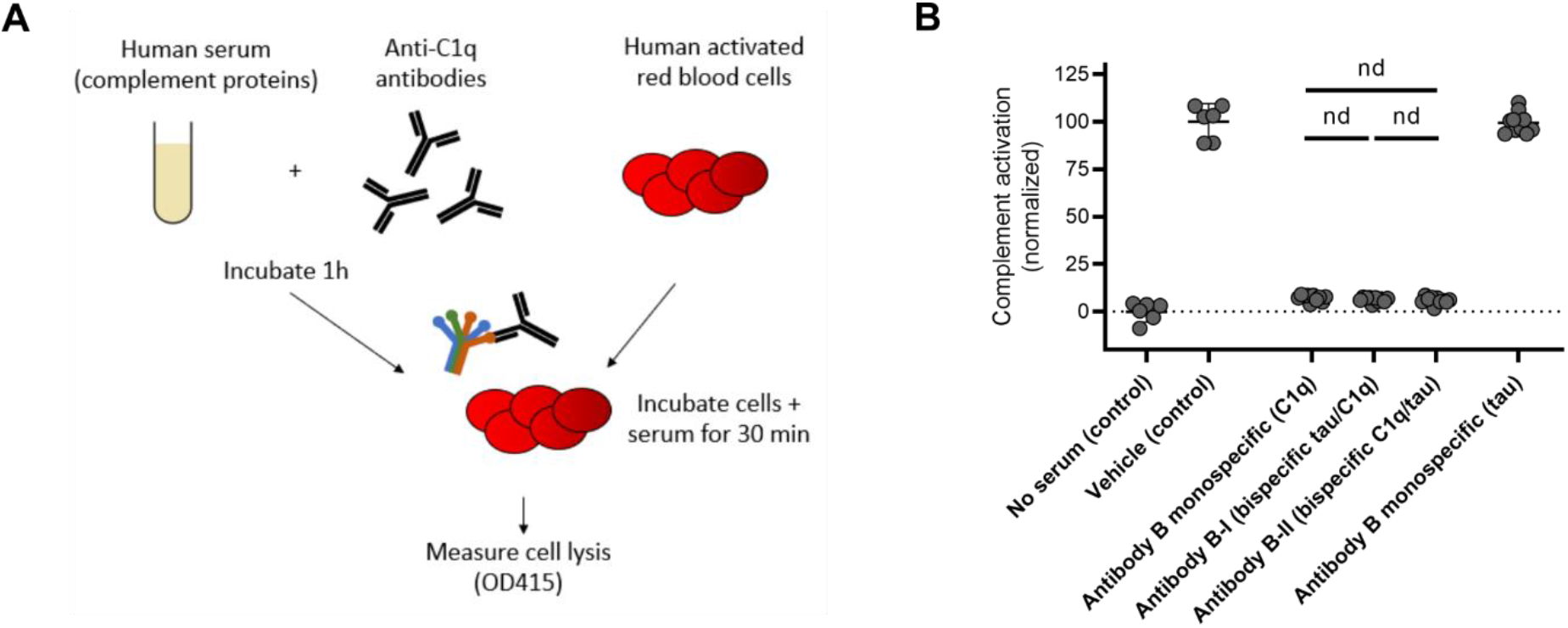
Comparison of mono- and bispecific antibodies in their ability to inhibit the classical complement pathway. (**A**) Schematic representation of the modified version of the CH50 assay to measure inhibition of classical complement activation. (**B**) Comparison of antibodies in the modified CH50 assay. Horizontal lines represent the mean and error bars represent the 95% confidence intervals. Only comparisons between monospecific antibodies and their bispecific counterparts are highlighted here. Detailed information is described in the corresponding results section.

### Inhibition of classical complement

C1q is the initiating factor of the classical complement pathway, which ultimately culminates into the lysis of the cells *via* formation of the membrane attack complex [78]. Classical complement activation can be initiated after antibodies bind to their targets (e.g. bacteria). C1q then binds to the Fc domain of IgG molecules to trigger the classical complement cascade [75]. The level of classical complement activation can be quantified using the widely used complement hemolysis 50% (CH50) assay [62]. In this assay, IgG-coated red blood cells are incubated with human serum – which contains all complement proteins. By making a plasma dilution curve and examining the resulting hemolysis, this assay can be used to estimate complement activity in the blood. This assay can be modified to test the neutralizing effect of C1q antibodies, which is accomplished by pre-incubation of serum with the experimental antibodies before application to the red blood cells [33] (Figure 5A).

We used this assay to compare the complement neutralizing effects of Antibody B_mono-C1q_, Antibody B-I and Antibody B-II (Figure 5B). A condition without serum was used a negative control to estimate 0% classical complement activation. Serum without experimental IgG was used as a positive control to estimate 100% classical complement inhibition. We used a dose that was previously reported to lead to full neutralization of classical complement [33]. Indeed, Antibody B_mono-C1q_ inhibited classical complement activity to a mean of 7.2% (corrected p= <0.0001, CI 6.0–8.5). As expected, antibody B_mono-tau_ which does not bind C1q, was completely inactive in this assay (corrected p= 0.882; mean value of p=99.39, CI 95.5– 103.3). This demonstrates that the observed effect could be explained by selective neutralization of C1q. Antibody B-I potently neutralized classical complement (corrected p= <0.001, mean value of 6.4%, CI 5.4– 7.3), as did Antibody B-II (corrected p= <0.001, mean value of 5.9%, CI 4.6–7.3) No differences were observed between antibody B_mono-tau_ and Antibody B-I (corrected p= 0.218) or Antibody B-II (corrected p= 0.116). Likewise, no differences were observed between Antibody B-I and Antibody B-II (corrected p= 0.546).

## Discussion

This study describes the development and use of bispecific antibodies as a new approach to simultaneously target multiple pathological targets. For this proof-of-principle study we selected αSyn and C1q because tau pathology often co-occurs with Lewy body pathology as well as classical complement activation. However, this approach can in principle be extended to any desired combination of targets. The affinities of the antibodies developed here range from comparable to one order magnitude lower in comparison to their monospecific counterparts. It is possible that antibodies may deviate slightly from the originally described versions because of the different IgG backbone and bispecific format. Importantly, the bispecific antibodies retained their ability to inhibit tau aggregation, cellular tau seeding and classical complement-mediated hemolysis. This was not dependent on the location of the target binding CDRs on the antibody.

Several other interesting approaches have been described to simultaneously target multiple targets. Vaccines that in parallel target Aβ plus tau [18], αSyn [52] or complement protein C5a [11] showed efficacy in transgenic animals. Although this is a highly promising approach, it remains to be determined to what extent vaccines are a suitable tool for treating neurodegenerative disorders. The main limitation is the lack of control over antibody titers – which may be particularly problematic in the context of aging [53]. In contrast to monoclonal antibodies, it is not clear that all antibodies raised naturally by the vaccine will have acceptable affinity and therapeutic efficacy. Furthermore, it is not possible to change the effector function of the resulting antibodies, possibly leading to undesirable and even irreversible neuroinflammation [47]. Finally, it is unclear to what extent monoclonal antibodies enter the brain parenchyma. These limitations led to the rise of engineered monoclonal antibodies with blood-brain barrier shuttles [35, 41], which is not possible with naturally produced antibodies in response to a vaccine. Bispecific antibodies, like the ones used in this study, can be engineered to be effector-neutral and have increased brain uptake.

Several studies described antibodies that bind to the β-sheet–rich structures, of which NPT008 is currently undergoing clinical trials [27, 28, 31, 48, 56]. This is a fascinating approach with the potential to simultaneously target multiple amyloidic proteins. It is, however, still an open question whether conformation-selective antibodies bind the full range of potential pathogenic states of molecules: misfolded monomers, soluble oligomers, protofibrils, and fibrils. In addition, this would only work as a combination therapy against aggregated proteins. With the advent of promising neuroinflammation-related targets for the treatment of neurodegenerative disorders, this might pose a potential limitation [24, 78]. However, since these antibodies can also be developed in bispecific formats, this opens the door to developing antibodies that can bind to multiple aggregated proteins with one binding site and neuroinflammation-related target with another.

An interesting recent study describes the functional characterization of a small molecule that can simultaneously target monomeric tau and αSyn [25]. In addition, an oligomer-specific small molecule anle138b reduces both αSyn and tau aggregation *in vitro* and reduces both αSyn and tau pathology in mouse models [9, 20, 49, 80, 81]. The major advantage of small molecules compared to monoclonal antibodies is the low cost. However, in contrast to the high specificity of therapeutic monoclonal antibodies, it is unclear to what extent bispecific small molecules have low off-target binding. Furthermore, this approach is mostly likely not feasible for any combination of targets. In contrast, the modular nature of bispecific antibodies makes it easy to construct them against a wide range of target combinations.

### Limitations

The main limitation of this study is that we focused only on a single bispecific antibody format. It is therefore unclear how these results translate to other multispecific antibody formats. Furthermore, Antibody B-I and B-II had approximately one order of magnitude lower affinity to tau. This suggests that further optimization of these bispecific antibodies is required to retain the affinity of the monospecific parent antibodies.

### Conclusion

In conclusion, we present a previously uncharacterized approach to simultaneously target multiple targets involved in pathological processes in neurodegeneration. The concept of this type of bispecific antibody expands the toolkit with treatment options for neurodegenerative disorders. Importantly, when only one of the two targets is present, the bispecific antibody will function just like a regular monospecific antibody. This demonstrates that the additional binding capacity does not come at the cost of decreased overall antibody functionality compared to the original antibody. A wide range of multispecific antibody formats have been described in the literature, which may each have their unique strengths when targeting different neurodegeneration-related targets [83]. Bispecific antibodies are therefore a promising approach and could be explored against a wide range of target combinations to obtain potential synergistic effects.

## Abbreviations

αSyn: Alpha-synuclein
Aβ: Amyloid beta
AD: Alzheimer’s disease
BLI: Bio-Layer Interferometry
CDR: Complementarity-determining region
DLB: Dementia with Lewy bodies
Fc: Fragment crystallizable
FRET: Fluorescence resonance energy transfer
IgG: Immunoglobulin G
NFT: Neurofibrillary tangle
PD: Parkinson’s disease
RT: Room temperature
scFv: Single-chain variable fragment

## Ethical Approval and Consent to participate

The brain samples were obtained from The Netherlands Brain Bank (NBB), Netherlands Institute for Neuroscience, Amsterdam (open access: www.brainbank.nl). All Material has been collected from donors for or from whom a written informed consent for a brain autopsy and the use of the material and clinical information for research purposes had been obtained by the NBB. The project has been reviewed and approved by the ethical board of the NBB.

## Authors’ information

Wim Hendricus Quint

Irena Matečko-Burmann

Irene Schilcher

Tina Löffler

Michael Schöll

Björn Marcus Burmann

Thomas Vogels

## Authors’ contributions

WHQ, IMB, IS, TL and TV performed experiments and analyzed data. TV drafted the manuscript. MS and BMB provided resources and scientific input. All authors contributed to writing of the manuscript and approved the final version.

## Funding

This study was funded by Maptimmune.

## Acknowledgements

We would like to thank the patients and their families for their donations and the Netherlands Brain Bank for providing the material for this project.

## Availability of data and materials

Data and materials are available from the corresponding author upon reasonable request.

## Competing interests

TV & WQ are major shareholders of Maptimmune BV, which commercially develops treatment and diagnostics for neurodegenerative disorders (unrelated to the current publication). IS & TL are employees of QPS Austria, which performed contract research for this research project. All other authors report no conflicts of interest.

